# The Heat shock protein 70 machinery is crucial in the production of infectious chikungunya virus progeny

**DOI:** 10.64898/2026.01.26.701664

**Authors:** M. van der Laan, S. Verwimp, O.T. Johnson, E.M. Bouma, K. Trappeniers, F.E. Visscher, H.H. van den Ende-Metselaar, D.P.I. van de Pol, L. Delang, J.E. Gestwicki, H.H. Kampinga, J.M. Smit

## Abstract

Over the past decades, chikungunya virus (CHIKV), a re-emerging arthropod-borne alphavirus, has caused outbreaks in many (sub)tropical regions, but also in more temperate regions of the world, including Europe. CHIKV poses a significant health burden due to high infection rates during epidemics, symptom progression into chronic arthritic manifestations, and the lack of specific antiviral treatments. Multiple studies have shown that several viruses rely on host molecular chaperones, particularly the central Heat shock protein 70 (Hsp70), for replication. Hsp70s are guided by co-chaperones that drive their functionality and specificity. Here, we used chemical inhibitors of Hsp70-co-chaperone interactions to study the role of this molecular chaperone machinery in CHIKV replication. Our findings revealed that Hsp70 inhibition significantly reduces the CHIKV infectious particle production without affecting host-cell viability. Inhibition of the Hsp70-co-chaperone interaction primarily impedes the post-RNA replication stages of the CHIKV infectious cycle, affecting viral protein expression and reducing both the number and infectivity of released virions. Moreover, Hsp70 inhibition displayed antiviral activity in skin explants. Together, these results suggest that targeting the Hsp70 network could be a viable antiviral strategy against CHIKV infections.

**Author summary:** As obligatory intracellular parasites, viruses rely entirely on the machinery of the host cell to produce new viral particles. In our study, we investigated whether a specific group of host proteins, known as molecular chaperones, is important for the replication of chikungunya virus (CHIKV), a reemerging mosquito-borne virus that can cause long-lasting joint pains and lacks specific antiviral treatments. We focused on one key family of molecular chaperones, called Heat shock protein 70s (Hsp70s), which supports protein folding and quality control in cells. We used chemical compounds to block Hsp70 function and observed that CHIKV replication was strongly reduced, while host cells remained healthy. We found that Hsp70 is especially important in the later stages of the virus life cycle, where it helps produce viral proteins and new infectious virus particles. When Hsp70 was blocked, fewer and less infectious virus particles were produced. We also showed that this effect holds true in biopsies of mouse skin tissue, which mimics the initial site of infection. These findings identify Hsp70 as an important host factor for CHIKV that may serve as a potential target for new antiviral therapies.

## Introduction

Chikungunya virus (CHIKV) is an alphavirus within the *Togaviridae* family, primarily transmitted to humans by mosquitoes, making it a member of the genetically diverse group of arthropod-borne viruses (arboviruses)[1,2]. Over the past decades, CHIKV has re-emerged explosively, causing millions of infections across Central America[3], and even sporadic outbreaks in Europe[4–6]. Approximately 85% of individuals infected with CHIKV develop chikungunya fever, which is characterized by symptoms such as fever, rash, headache, and myalgia[1,7]. The morbidity and economic burden[8] in affected regions is particularly severe, as over 40% of chikungunya fever cases progress to chronic (poly)arthralgia and/or polyarthritis, which can persist for months or even years after the initial infection[9]. Despite extensive efforts[10,11], there are currently no specific antiviral treatments available for either acute or chronic CHIKV-induced disease symptoms.

CHIKV infection is initiated by the attachment of the virion to the host cell via the viral E1 and E2 transmembrane glycoproteins[12–14]. Next, the virion is internalized via clathrin-mediated endocytosis, whereafter the viral membrane fuses with the membrane of the early endosome[15,16], and the positive-sense single-stranded RNA genome is released into the cytoplasm. The released viral RNA (vRNA) includes two coding regions: the first region encodes the four non-structural proteins (nsPs) that form the viral replication complex (RC), and the second and subgenomic region encodes the structural polyprotein essential for particle assembly[1,17]. vRNA replication takes place in membrane-derived spherules that contain a full-length negative-sense RNA template and the RC formed from nsPs[18,19]. From this template, the genomic vRNA and subgenomic vRNA are transcribed. Next, the structural polyprotein (C-E3-E2-6K-E1) is translated, and the capsid protein is autoproteolytically cleaved from the polyprotein in the cytoplasm. Thereafter, the remaining polyprotein is translocated to the ER, where the glycoproteins E1 and E2 are processed and folded. The glycoproteins mature while passing through the Golgi network and are eventually expressed at the plasma membrane of the infected cells[17]. The capsid proteins interact with the genomic vRNA to assemble a nucleocapsid, and the subsequent interaction of the formed nucleocapsid with the cytoplasmic tail of the E2 glycoprotein, which is expressed at the plasma membrane, initiates the assembly and budding of a progeny virion[20].

Viruses only carry what is minimally essential in their genome and are therefore highly dependent on the host cell for several processes, such as the formation of functional viral proteins[21]. To support the production of viral proteins needed for RNA replication and the assembly of progeny virions, viruses hijack the host cellular folding and assembly systems[22]. Central components of the intracellular protein quality control network are the ubiquitous molecular chaperones of the Heat shock 70kDa protein (Hsp70) family. Each major cellular compartment therefore contains an Hsp70 homolog, which facilitates protein folding, degradation, transport across membranes, and the assembly and disassembly of multiprotein complexes[23]. Hsp70 homologs have a highly conserved structure and mode of action[24], functioning through ATP-driven cycles of binding and release of client proteins. This process minimally requires two co-chaperones: J-domain proteins (JDPs)[25] and nucleotide exchange factors (NEFs)[26]. JDPs are thought to first bind to the protein client and next recruit and position ATP-bound Hsp70 to the client, resulting in ATP hydrolysis, which causes a conformational change in Hsp70 that will stabilize the Hsp70-client interaction[25]. NEFs then bind to Hsp70 in its ADP-bound state and can cause ADP dissociation, resulting in reloading of Hsp70 with ATP, generally causing client release[26]. This ATP-driven cycle is repeated until the correct folding or assembly of the client is achieved. Most cellular compartments contain a specific subset of JDPs and NEFs that not only promote the functionality of Hsp70 but also drive the diversity and specificity of the machinery[23,27].

The involvement of the Hsp70 machines in various diseases, including cancer and neurodegeneration, has spurred research into developing chemical modifiers of their functions[28,29]. Given that co-chaperones drive the functionality and specificity of Hsp70, research on these chemical inhibitors soon shifted focus to disrupting the Hsp70-co-chaperone interaction. One of the earliest efforts led to the discovery of the benzothiazole-rhodacyanine compound MKT-077, first described by Wadhwa and colleagues[30]. MKT-077 binds a conserved allosteric site within the NBD of Hsp70, preventing NEF binding and thus stalling the ATPase cycle of Hsp70[31]. While MKT-077 demonstrated strong anti-tumor activity, it was also rapidly metabolized[32]. This limitation prompted a medicinal chemistry campaign aiming to enhance the potency[31] and pharmacokinetic properties of MKT-077[33], resulting in approximately 400 analogs of MKT-077 named JG-compounds. Although the binding site for these JG-compounds is conserved across the main members (HspA1, HspA8, HspA5, and HspA9) of the Hsp70 family[31], it was found that these compounds display differential localization patterns and varying activity against distinct diseases, such as cancer or viral infections[34]. While further research is required to fully understand these findings, they suggest that these compounds do not target all Hsp70 isoforms to the same extent, thereby selectively modulating a specific subset of processes mediated by Hsp70.

Replication of multiple viruses has been shown to depend on specific Hsp70-mediated processes, presenting an opportunity to target this chaperone machinery to reduce virus replication[22]. Targeting Hsp70 and its co-chaperones via short-hairpin RNAs (shRNAs) or allosteric inhibitors has previously been demonstrated to effectively reduce the replication of various viruses. For instance, the Endoplasmic Reticulum (ER)-resident Hsp70 (HspA5) and its co-chaperones control the ER exit of the nonenveloped DNA virus, Simian virus 40 (SV40)[35]. Additionally, allosteric inhibitors of Hsp70 were found to interfere with multiple steps of the replication cycle, including entry and vRNA replication, of the more closely related arboviruses, dengue virus (DENV)[36] and Zika virus (ZIKV)[37]. For DENV, specific JDP co-chaperones aid in the entry phase (DNAJC9, DNAJC16, DNAJC18), while others (DNAJA2, DNAJB6, DNAJB7, DNAJB11, and DNAJC10) assist in post-entry steps [36] of its replication cycle. Importantly, restricting viral replication using allosteric inhibitors of the Hsp70-co-chaperone interactions (JG18, JG40, or JG345) did not lead to viral resistance (DENV[36] and ZIKV[37]) and offered protection against virus-induced disease in mice (ZIKV[37]). These findings highlight the potential of chemically inhibiting Hsp70 and its co-chaperones to both study the role of this molecular chaperone network in viral infections and potentially even serve as a broad-spectrum therapeutic strategy against diverse viruses.

In analogy, we decided to investigate whether Hsp70 plays a role in CHIKV replication and if it could serve as a therapeutic strategy against CHIKV infections. Due to the differences in the subcellular location of CHIKV viral replication compared to previously explored systems, we tested multiple Hsp70 inhibitors (JG-compounds) and found that a subset of them significantly impaired CHIKV replication. Somewhat unexpectedly, inhibition was unrelated to the early stages of viral replication (i.e., cell entry and vRNA replication). Rather, Hsp70 seemed to play critical roles in the later stages of the CHIKV replication cycle. Specifically, we found that Hsp70 inhibition reduced viral protein expression and particle production, with an even stronger effect on the infectivity of the produced viral progeny. Our results indicate a critical (and differentiated) role for the molecular chaperone Hsp70 in the replication cycle of CHIKV.

## Materials and methods

### Cells

The human bone osteosarcoma epithelial cell line U2OS (ATCC HTB-96) and human foreskin fibroblast cell line HFF-1 (ATCC SCRC-1041) were maintained in Dulbecco’s minimal essential medium (DMEM), high glucose, GlutaMAX^tm,^ and sodium pyruvate (Gibco), supplemented with penicillin-streptomycin (P/S; 100U/mL, Gibco) and 10% or 15% fetal bovine serum (FBS, Lonza), respectively. Baby hamster kidney cells (BHK-21; ATCC CCL-10) were maintained in RPMI medium supplemented with 10% FBS, and P/S. BHK-21 cells clone 15 (BHK-15) were a kind gift of Richard Kuhn (Purdue University) and are not commercially available. Renal African Green monkey Vero-WHO cells were obtained from ATCC (ATCC CCL-81, Reference Cell Bank 10-87). BHK-15 and Vero-WHO cells were maintained in DMEM, high glucose, containing 10% or 5% FBS, respectively, and P/S. All cell lines tested negative for Mycoplasma and were cultured at 37°C and 5% CO^2^.

### Mice

In-house-bred AG129 mice (10–18 weeks old), which lack both IFN-α/β and IFN-γ receptors, were used for the preparation of mouse skin explants. Mice were housed under standard conditions in individually ventilated isolator cages (IsoCage N Biocontainment System, Tecniplast), maintained at 18–23°C with a 14 h:10 h light–dark cycle and 40–60% relative humidity. Cage enrichment was provided, and animals had ad libitum access to food and water. All animal care and experimental procedures were approved by the Ethical Committee of the University of Leuven (license M020/2020) and conducted in accordance with institutional guidelines and those of the Federation of European Laboratory Animal Science Associations (FELASA).

### Hsp70 inhibitors

Hsp70 inhibitors (JG-compounds) were synthesized as described previously[31,33] and were >95% pure by HPLC. JG18, JG40, JG98, JG231, JG345, and VER-155008 were all dissolved in dimethyl sulfoxide (Sigma-Aldrich) to a working stock of 5 mM and later diluted in culture media to the working concentration.

### MTS assay

Cytotoxicity of the Hsp70 inhibitors in U2OS and HFF-1 cells was assessed using the CellTiter® 96 AQ_ueous_ Non-Radioactive Cell Proliferation Assay (MTS assay, Promega). Cells were seeded in a 96-well plate and incubated with increasing concentrations of the Hsp70 inhibitors or the DMSO control at 37°C and 5% CO_2_. At 16 h post-treatment, 20 µL MTS/PMS solution was added, and cells were further incubated for 2 h. Subsequently, 25 µL 10% sodium dodecyl sulfate was added, and the absorbance was measured at 490 nm with a microplate reader (GloMAX discover, Promega). Data was expressed as a percentage of metabolically active cells relative to the corresponding DMSO control.

### Viruses and quantification

The infectious clone based on the CHIKV La Réunion (LR) strain 2006 OPY1 was kindly provided by prof. Andres Merits (University of Tartu, Estonia). Virus production was performed as described before[15]. Briefly, RNA was *in vitro*-transcribed and electroporated into BHK-21 cells to produce the CHIKV-LR strain. The CHIKV-S27 strain was kindly provided by prof. S. Guenther (Bernhard-Nocht-Institute for Tropical Medicine) and was isolated in 1953 in Tanzania. The CHIKV-99659 strain was a kind gift from prof. M. Diamond (Washington University, School of Medicine) and was isolated in 2014 in the British Virgin Islands in the Caribbean. The CHIKV Indian Ocean strain 899 (GenBank FJ959103.1), hereafter CHIKV-889, was a generous gift of Prof. Drosten (University of Bonn, Bonn, Germany)[38]. All CHIKV strains were propagated on Vero-WHO cells to produce working stocks, as previously described[39]. DENV-2 strain 16681 was propagated on C6/36 cells, as previously described[40]. The infectious titer in plaque-forming units (PFU) was determined via plaque assay on Vero-WHO (CHIKV) or BHK-15 (DENV) cells. Vero-WHO cells or BHK-15 cells were seeded in a 12-well plate at a density of 1.3x10^5^ or 9.0x10^4^ cells per well, respectively. At 24 h post-seeding, the cells were infected with 10-fold serial dilutions of the samples. At 1.5 hours post-infection (hpi) an overlay of 1% SeaPlaque Agarose (Lonza, Switzerland) prepared in MEM was added and plaques were counted at 46 hpi for CHIKV and 5 days post-infection (dpi) for DENV. The number of CHIKV genome-equivalent copies (GECs) was quantified by reverse transcriptase quantitative PCR (RT-qPCR) on a Biorad CFX (Bio-rad) using primers for the E1 sequence, as previously described[41]. The number of GECs was determined using a standard curve (R^2^ >0.995) of a quantified cDNA plasmid that contains the CHIKV E1 sequence (pCHIKV-LS3 1B).

### Antiviral assays

U2OS or HFF-1 cells were infected with CHIKV or DENV at the indicated multiplicity of infection (MOI). Virus inoculum was prepared in cell culture media containing 2% FBS to which Hsp70 inhibitors or the DMSO control were added. Infection was performed at 37°C and 5% CO^2^ for 1.5 h on a shaker. After incubation with the inoculum, cells were washed three times with unsupplemented DMEM. Thereafter, fresh culture media containing 10% FBS and 1.5% HEPES (Gibco) with compounds or the DMSO control were added. Supernatants were collected at 9 hpi (CHIKV) and 24 hpi (DENV), centrifuged to remove cell debris, and subjected to plaque assay or RT-qPCR as described above.

### Mouse Skin Explants

Skin explants were prepared from AG129 mice. The back of the mice was shaved as thoroughly as possible using an electric razor, and a large section of back skin was excised and placed in Dulbecco’s Modified Eagle Medium (DMEM, Gibco) supplemented with 10% FBS, 1% P/S, and 0.1% gentamicin (Gibco). The tissue was kept on ice until further processing. Subcutaneous adipose tissue was carefully removed using forceps and scissors, and the skin was then cut into ∼1 cm² pieces. Following a PBS wash, individual skin pieces were placed epidermis-side up in a 24-well plate (Falcon) containing 1 mL of the supplemented DMEM per well. Samples were incubated at 37°C and 5% CO_2._ To evaluate the *ex vivo* antiviral efficacy of JG231, skin explants were infected with CHIKV-899 (10^6^ PFU/ml) and treated with 2 µM of compound or an equal volume of DMSO per skin explant in a total volume of 1 mL culture media supplemented with 2% FBS. Skin samples were incubated for 1.5 h, after which the medium was removed, and skin explants were washed three times with PBS. Subsequently, skin explants were incubated with 1 mL of fresh supplemented DMEM medium, to which compounds were again added at similar concentrations, followed by incubation at 37°C. The skin explants were harvested at 2 dpi, and GECs were measured by RT-qPCR and infectious titers were defined by end-point titrations.

### GEC quantification in skin explants

To quantify GECs in skin explants, tissues were first homogenized in Precellys tubes containing 2.8 mm zirconium oxide beads (Bertin Instruments) and TRK lysis buffer (Omega-Biotek). Homogenization was performed using an automated homogenizer (Precellys24, Bertin Instruments) at 7600 rpm for 20 seconds, for two cycles with a 20-second interval. Tissue homogenates were centrifuged at 15,000 rpm for 10 minutes at 4°C. RNA was isolated from the supernatants using the E.Z.N.A. Total RNA Kit I (Omega-Biotek), following the manufacturer’s protocols. Quantification of viral RNA was performed using quantitative reverse transcription PCR (qRT-PCR) in a 25 µl reaction volume containing: 13.94 µL nuclease-free water (Promega), 6.25 µL master mix (Eurogentec), 0.375 µL each of forward (5′-CCG ACT CAA CCA TCC TGG AT-3′) and reverse (5′-GGC AGA CGC AGT GGT ACT TCC T-3′) primers at 150 nM final concentration (IDT), 1 µL of probe (5′-FAM-TCC GAC ATC ATC CTC CTT GCT GGC-TAMRA-3′; 400 nM final; IDT), 0.0625 µL reverse transcriptase (Eurogentec), and 3 µL of RNA template. Amplification was performed on a QuantStudio 5 Real-Time PCR system (ThermoFisher Scientific) with the following program: 30 minutes at 48°C, 10 minutes at 95°C, followed by 40 cycles of 15 seconds at 95°C and 1 minute at 60°C. A standard curve was generated from a ten-fold dilution series of CHIKV reference plasmid DNA. The limit of quantification (LOQ) was defined as the lowest viral load detectable in 95% of replicates, considering tissue weights and buffer volumes.

### End-point titrations of skin explants

Infectious virus levels in skin explants were quantified by end-point titration on Vero cells. Cells were seeded in 96-well plates at 10⁴ cells/well and incubated overnight at 37°C with 5% CO_2_. Tissue homogenates were prepared in Precellys tubes filled with culture media supplemented with 2% as described above, centrifuged, and ten-fold serial dilutions of the supernatants were added to the cells in triplicate. At 3 dpi, cells were microscopically scored for CHIKV-induced cytopathogenic effects (CPE) relative to uninfected controls. Viral titers were calculated as TCID₅₀/mL (tissue culture infectious dose 50%) using the Reed and Muench method[42]. The limit of quantification was defined as the lowest titer that could be reliably measured, accounting for tissue input and dilution factors.

### Quantification of intracellular vRNA copies

U2OS cells were infected with CHIKV at MOI 10 with or without JG98, as described above. At 4, 6, and 8 hpi cells were washed with PBS and intracellular RNA was extracted using the RNAeasy Mini Kit (Qiagen) as per the instructions of the manufacturer. Intracellular vRNA copies were determined via independent RT-qPCR reactions using primers targeting the E1 or nsP1 sequence, as described before[43].

### Analysis of viral protein expression by Western-Blotting

To determine the effect of Hsp70 inhibition on expression levels of viral proteins, U2OS cells were infected with CHIKV at MOI 10 in the presence or absence of JG98, JG345, or the DMSO control. At 9 hpi cells were washed with PBS and protein extraction was performed using the RIPA lysis buffer system (Santa Cruz) containing protease inhibitors. Briefly, 12-well plates with 2.0x10^5^ U2OS cells per well were kept on ice, and ice-cold RIPA buffer was added. After 10 minutes of incubation, cells were scraped, and the cell lysate was transferred to a tube. Cell lysates were then incubated on ice for 20 minutes and vortexed at 10-minute intervals. Samples were diluted in 4x SDS sample buffer (Merck) and heated at 95°C for 5 min before separation by SDS-PAGE. Separated proteins were transferred onto PVDF membranes using the Trans-Blot Turbo system (Bio-Rad). The primary antibodies used were anti-vinculin (1:1000, mouse monoclonal, hVIN1, Merck), anti-nsP2 (1:1000, mouse monoclonal, ABM3F3.2E10, Abgenex), anti-capsid (1:1000, rabbit polyclonal, kindly provided by G. Pijlman (Wageningen University, the Netherlands)), and anti-E1 (1:1000, rabbit polyclonal, kindly provided by G. Pijlman). The secondary, HRP-conjugated antibodies, donkey anti-rabbit HRP (1:5000, Alpha Diagnostic International), and goat anti-mouse HRP (1:2000, A8924, Merck), were used. Protein signal intensities were quantified using the ImageQuant TL8.1 software.

### Flow cytometric analysis of viral protein expression

To determine the expression levels of viral proteins in CHIKV-infected cells, U2OS cells were infected with CHIKV at MOI 10 with or without treatment with JG98 and JG345. At 9 hpi, cells were washed with PBS, trypsinized and/or scraped, and fixed with 4% paraformaldehyde (Alfa Aeser). For intracellular staining for E2 and capsid, fixed cells were permeabilized before staining them with the mouse anti-capsid antibody (1:1000, the Native Antigen Company) and the Alexa Fluor 647-conjugated rabbit anti-mouse antibody (1:2000, Life Technologies, A-21443), the Alexa Fluor 647-labeled anti-E2 antibody (1:1000, 6A11, Novus Biologicals), or the Alexa Fluor 647-labeled mouse IgG2b isotype control ((1:1000, IC0041R, R&D systems). Flow cytometry was performed using the NovoCyte Quanteon (Agilent) and analyzed using FlowJo vX.07. A schematic overview of the gating strategy can be found in **Fig. S2**.

### Time-of-drug-addition experiment

U2OS cells were treated with Hsp70 inhibitors at the indicated concentrations before, during, or after the incubation with the virus inoculum (MOI 1). A schematic representation of the different time points can be found in **Fig. 4A**. For the conditions that required pre-incubation (conditions 1-3), cells were treated with the compounds for 2 h and washed 3 times with unsupplemented DMEM before the virus inoculum in DMEM with 2% FBS and with or without compounds was added. After 1.5 h, the inoculum was removed, and cells were washed 3 times with unsupplemented DMEM before DMEM with 10% FBS and 1.5% HEPES was added. For condition 4, the compounds were only present during the incubation with the virus inoculum. For condition 5, the compounds were re-added to the cells after the inoculum was removed. For the last conditions (6-8), the inhibitors were added after the removal of the inoculum at 1.5, 4, and 6 hpi. Supernatants were collected at 9 hpi, and cell debris was removed by centrifugation. Viral titers in the supernatant were determined via plaque assay.

### Statistical analysis

All data sets were analyzed using the GraphPad Prism 10 software and are presented as mean±SEM. Statistical differences between conditions were assessed using One-Way ANOVA or Student T-test and were presented when p≤0.05 with *p ≤ 0.05, **p ≤ 0.01, ***p ≤ 0.001, ****p ≤ 0.001.

## Results

### Inhibition of the Hsp70 machinery reduces the production of CHIKV infectious particles

To assess the involvement of the Hsp70 network in the CHIKV replication cycle, we selected four Hsp70 inhibitors (JG18, JG40, JG98, and JG345) from the medicinal chemistry campaign encompassing ∼400 MKT-077 analogs[31,33] based on their reported antiviral activity against multiple flaviviruses[36,37,44]. The effect of these compounds was evaluated in human bone osteosarcoma (U2OS) cells, which are known to be both susceptible and permissive to CHIKV infection. First, we assessed the potential cellular toxicity of these compounds in U2OS cells using the MTS assay **(Fig. 1A)**. JG18 and JG40 exhibited minimal effects on cellular metabolic activity at concentrations up to 40 µM and 20 µM, respectively. To minimize compound use, the highest concentration used in subsequent experiments was set at 10 µM for both JG18 and JG40. JG98 and JG345 showed a stronger dose-dependent effect on cellular metabolic activity; therefore, the highest non-toxic dose (≥90% metabolically active cells) used in further experiments was 2 µM and 1 µM, respectively.

**Figure 1.**
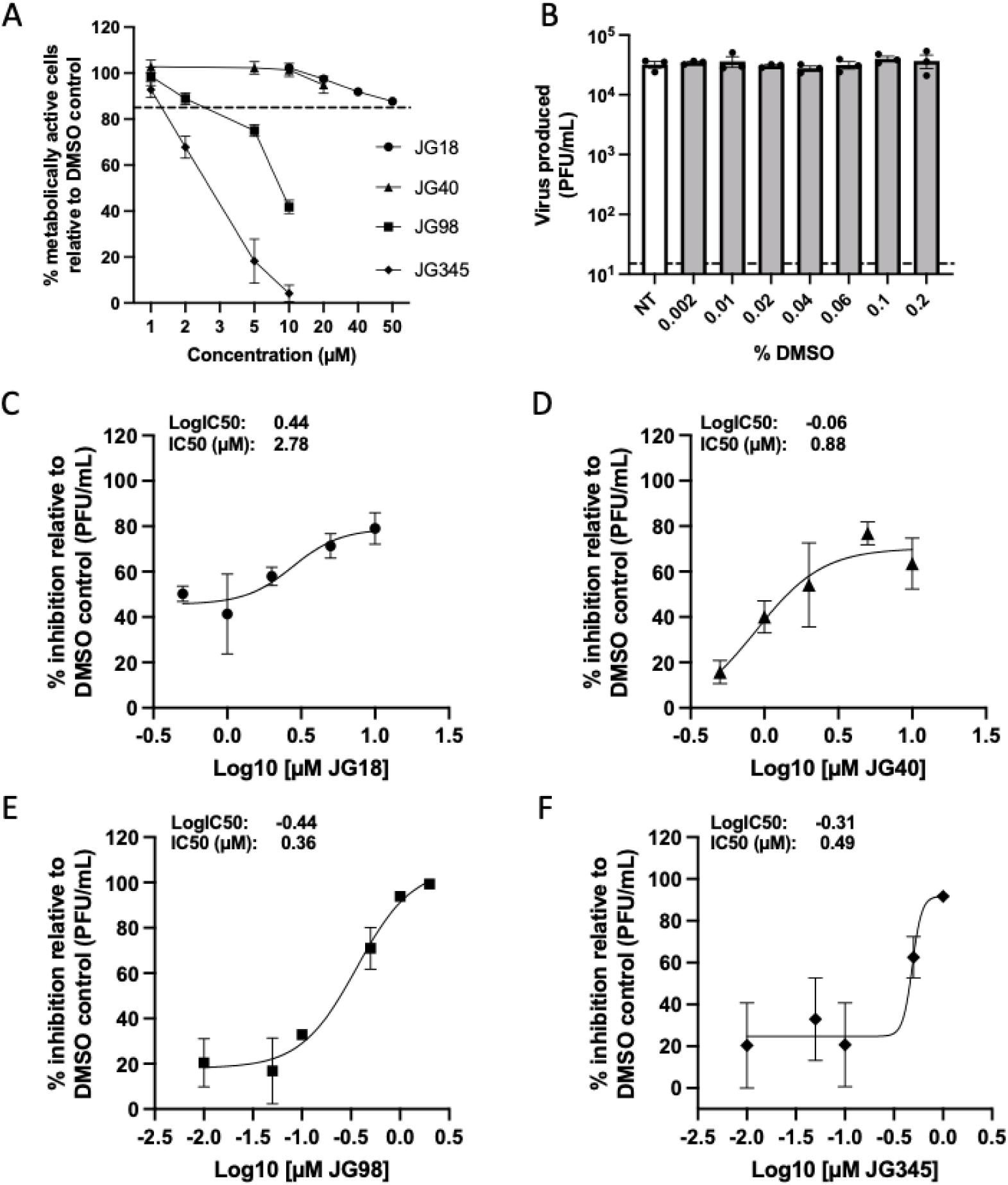
Hsp70 inhibitors reduce CHIKV progeny production at non-toxic concentrations. **(A)** U2OS cells were treated with increasing concentrations of the Hsp70 inhibitors or the equivalent volume of the DMSO control. The metabolic activity of the cells was assessed after 16 hours (h) of treatment using the MTS assay and was normalized to the DMSO control. A decrease of 15% in metabolically active cells was considered non-toxic (dotted line represents 85%). **(B-F)** U2OS cells were infected with CHIKV-LR OPY1 at multiplicity of infection (MOI) 1 in the presence of increasing concentrations of the **(B)** DMSO control (0.002% matches 0.1 µM JG-compound, 0.01% DMSO matches 0.5 µM JG-compound, etc.) or the Hsp70 inhibitors concentrations of **(C)** JG18, **(D)** JG40, **(E)** JG98, **(F)** JG345. Supernatants were collected at 9 hpi, and the number of infectious CHIKV particles was quantified using plaque assay on Vero-WHO cells. **(B-F)** Percentage inhibition was determined relative to the DMSO control, and IC50 (which corresponds to a 50% reduction in viral titer) per Hsp70 inhibitor was determined via non-linear regression. Data are presented as mean±SEM from three independent experiments. **(B)** Statistical differences were determined using One-way ANOVA and were presented when p≤0.05 with *p ≤ 0.05, **p ≤ 0.01, ***p ≤ 0.001, ****p ≤ 0.001.

Next, we assessed the impact of the JG-compounds on the CHIKV infectious particle production. To this end, U2OS cells were infected with CHIKV LR2006 OPY1 (CHIKV-LR), a clinical isolate derived from a patient on the La Réunion Island in the Indian Ocean. Infections were conducted in the presence or absence of the JG-compounds, or with the equivalent volume of the solvent DMSO. In the non-treated (NT) control and under increasing DMSO concentrations (max. concentration below 0.2%), on average 3.2x10^4^ and 3.5x10^4^ infectious virus particles were produced at 9 hours post infection (hpi) (**Fig. 1B**), demonstrating robust virus production and confirming that the solvent (DMSO) did not influence the production of infectious particles. With JG18, JG40, JG98, and JG345 treatment, we observed a dose-dependent reduction in the infectious particle production (**Fig. 1C-F**).

The IC50 values, representing a 50% reduction in viral titer, were calculated using non-linear regression (**Fig. 1C-F**, insets). Based on their IC50 values, JG98 (0.36 µM) and JG345 (0.49 µM) were found to be 1.8-7.7 times more effective against CHIKV-LR compared to JG18 (2.8 µM) and JG40 (0.9 µM). Additionally, at the highest tolerable dose, the maximal reduction in CHIKV-LR progeny production was lower with JG18 and JG40 (79.0% and 63.5%, respectively) than with JG98 and JG345 (99.3% and 91.7%, respectively). The substantial inhibition of the CHIKV-LR infectious progeny production by these distinct Hsp70 inhibitors suggests a critical role for Hsp70 in its replication cycle.

### The antiviral activity of Hsp70 inhibitors extends to different CHIKV strains

To rule out that our compounds exert strain-specific antiviral effects sometimes observed for antivirals[45,46], we tested the impact of JG18, JG40, JG98, and JG345 on two additional CHIKV strains from different genotypes. Specifically, we used the African S27 strain (CHIKV-S27), originally isolated in 1953[47], and the Caribbean 99659 strain (CHIKV-99659) from the Asian genotype, isolated in 2014[48]. In the non-treated and DMSO control groups, infection with the CHIKV-S27 strain resulted in the production of 2.6x10^5^ and 2.4x10^5^ infectious particles, respectively (**Fig. 2A**). For the CHIKV-99649 strain, 2.6x10^4^ and 2.3x10^4^ infectious particles were produced in the non-treated and DMSO control groups, respectively (**Fig. 2B**). These results confirmed that the solvent DMSO did not influence the production of infectious particles in either of the CHIKV strains tested. Consistent with our findings for the CHIKV-LR strain, treatment with JG98 and JG345 resulted in strong reductions in particle production for both CHIKV-S27 and CHIKV-99659 **(Fig. 2A and B),** whereas JG40 had only a moderate and JG18 had no significant effect on particle production in these strains. These results suggest that the antiviral effects of these Hsp70 inhibitors are consistent across different CHIKV strains, indicating a strain-independent mechanism of action.

**Figure 2.**
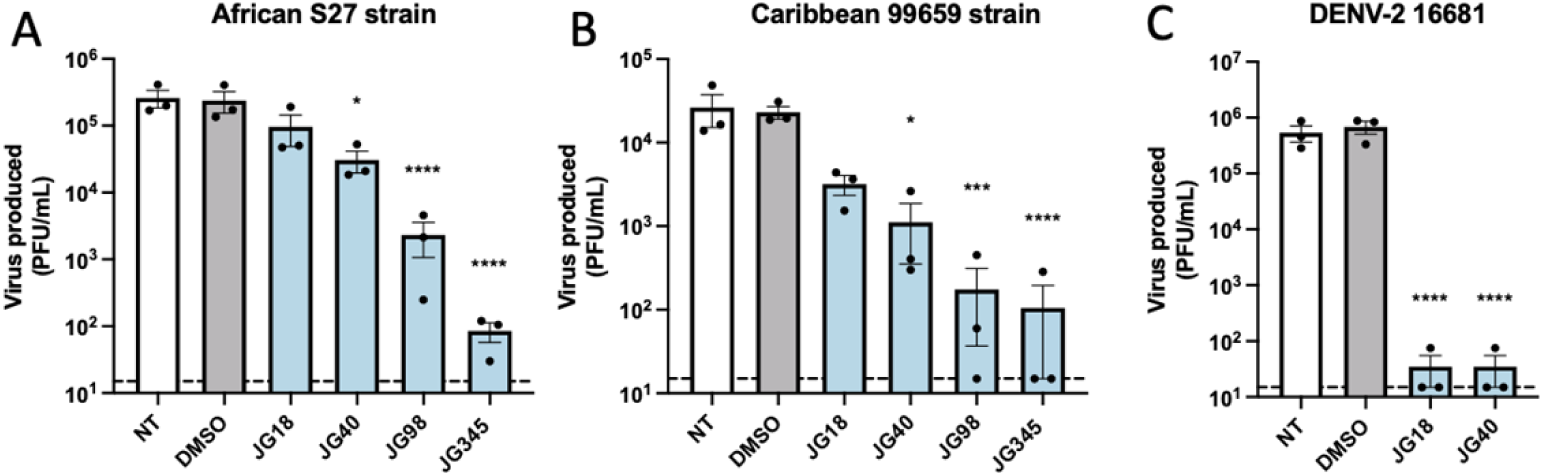
The antiviral activity of JG40, JG98, and JG345 extends to different CHIKV strains. U2OS cells were infected with the **(A)** African S27 CHIKV strain or the **(B)** Caribbean 99659 strain at MOI 1 and simultaneously treated with 10 µM, 10 µM, 2 µM, or 1 µM of JG18, JG40, JG98, or JG345, respectively, or the DMSO control. Production of infectious virus particles was assessed at 9 hpi. **(C)** U2OS cells were infected with DENV A2 (16681) at MOI 0.5 and treated with 10 µM JG18 or JG40, or the vehicle control. 24 hpi, supernatants were collected and DENV progeny production was analyzed via plaque assay on BHK-21 cells. Data are presented as mean±SEM from three independent experiments, and statistical differences were determined via one-way ANOVA and presented when p≤0.05 with *p ≤ 0.05, **p ≤ 0.01, ***p ≤ 0.001, ****p ≤ 0.001.

Interestingly, the drug efficacies observed for CHIKV differed from those reported for DENV, where JG18 and JG40 were much more effective in reducing viral replication[36]. To confirm the activity of JG18 and JG40, we assessed their inhibitory effects on DENV as a positive control and observed that they were highly potent in reducing the production of infectious DENV-2 particles (**Fig. 2C**). This not only confirms the high activity of JG18 and JG40 but also suggests that CHIKV and DENV-2 may rely on different Hsp70-co-chaperone interactions or interactions in distinct subcellular locations[34].

### Hsp70 inhibition reduces CHIKV progeny production in human skin fibroblasts

Given their superior efficacy against CHIKV in U2OS cells and their improved pharmacokinetics compared to JG18 and JG40[34], we decided to prioritize JG98 and JG345 for further investigation. To build on these results, we evaluated the antiviral activity of these compounds in human skin fibroblasts (HFF-1). The Hsp70 inhibitors displayed minimal cytotoxic effects on HFF-1 cells (**Fig. 3A)** and reduced the CHIKV progeny production to the limit of detection of the plaque assay (**Fig. 3B**).

**Figure 3:**
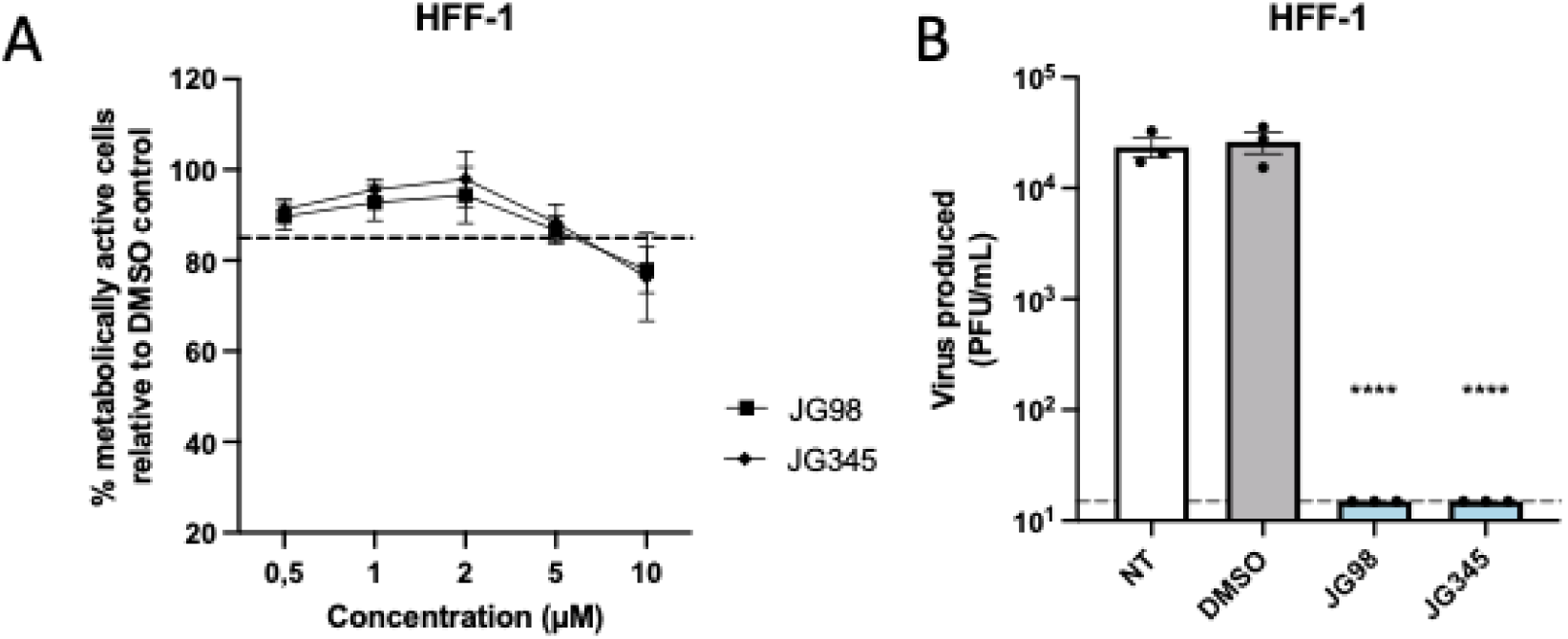
Hsp70 inhibition reduces CHIKV particle production in HFF-1 cells at non-toxic concentrations. **(A)** Metabolic activity of the HFF-1 cells was assessed after 16 h of treatment with increasing concentrations of the Hsp70 inhibitors or the equivalent volume of the DMSO control. The percentage of metabolically active cells is presented relative to the DMSO control. A decrease of 15% in metabolically active cells was considered non-toxic (dotted line represents 85%). **(B)** HFF-1 cells were infected with CHIKV-LR OPY1 at MOI 5 while treated with 2 µM or 1 µM of JG98 or JG345, respectively, and virus production was assessed at 9 hpi via plaque assay. Data are presented as mean±SEM from three independent experiments, and statistical differences were determined via One-way ANOVA and presented when p≤0.05 with *p ≤ 0.05, **p ≤ 0.01, ***p ≤ 0.001, ****p ≤ 0.001.

### Inhibitors of the Hsp70 network reduce the CHIKV particle production even when added late during infection

Hsp70 could play roles at multiple stages of the CHIKV lifecycle. To determine the specific stage of the CHIKV replication cycle that is affected by Hsp70 inhibition, we conducted a time-of-addition experiment (**Fig. 4A**). Both JG98 (**Fig. 4B**) and JG345 (**Fig. 4C**) significantly reduced particle production when present throughout the entire experiment (condition 1) or during the complete infection cycle (condition 5). Hsp70 inhibition during the early stages of CHIKV infection only (pre-entry and entry, conditions 2-4) resulted in minimal effects on viral progeny production. Inversely, when adding the compounds only during the later stages of CHIKV infection, specifically replication, translation, assembly, and budding (conditions 6-8), we still observed a strong, although not complete, inhibitory effect on infectious particle production for both JG98 (**Fig. 4B**) and JG345 (**Fig. 4C**). Remarkably, even when the inhibitors were applied only during the final three hours of infection (condition 8), there was a significant reduction in infectious particle production. Overall, these findings suggest that Hsp70 inhibition primarily affects the later stages of CHIKV infection, indicating a crucial role for Hsp70 in post-entry processes, such as RNA replication, protein translation, particle assembly, and/or budding.

**Figure 4:**
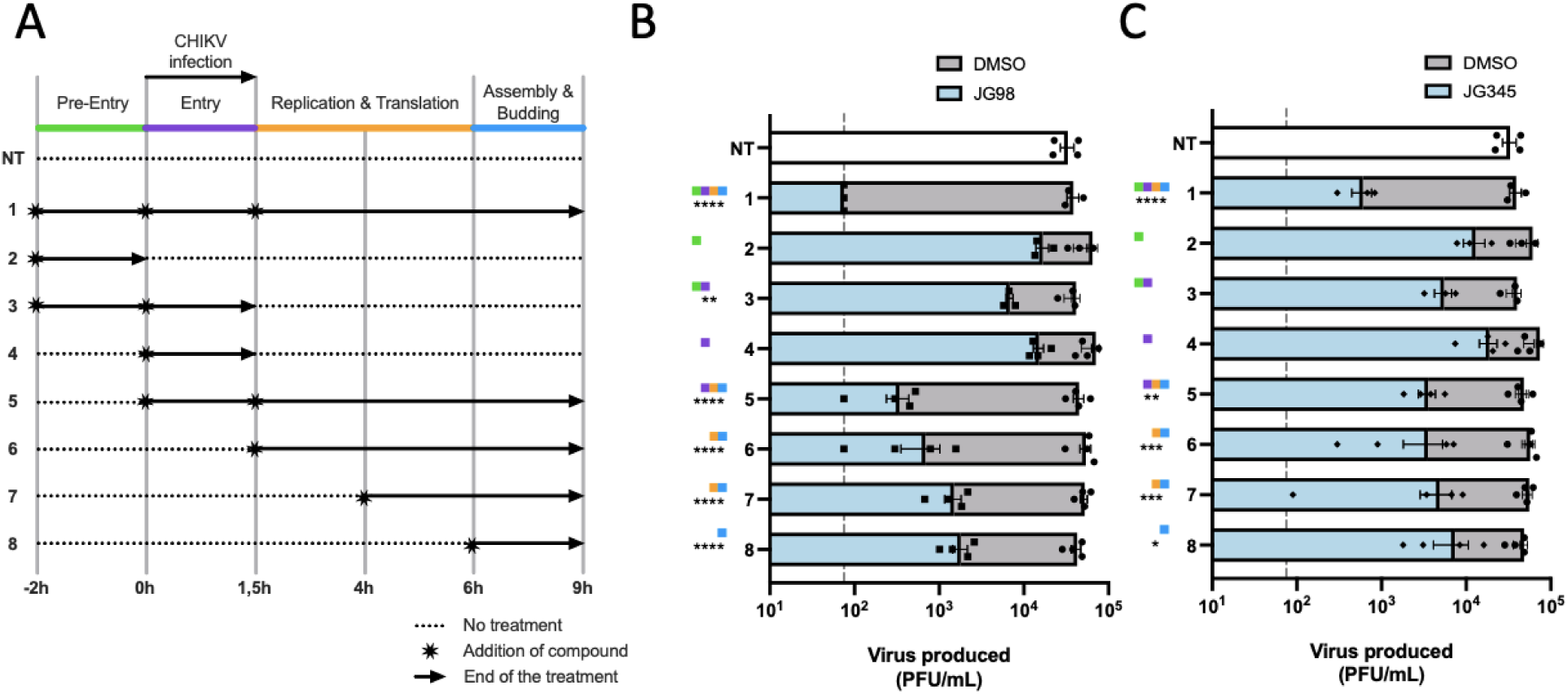
Hsp70 inhibitors are still effective when added during late steps in the CHIKV replication cycle. **(A)** Schematic representation of the time-of-drug-addition assay. U2OS cells were treated with **(B)** 2 µM JG98 or **(C)** 1 µM JG345 at the indicated time points (conditions 1-8) or with the DMSO control. Virus inoculum (CHIKV-LR at MOI 1) was present for 1.5 h, after which the inoculum was removed, and fresh medium was added with or without the Hsp70 inhibitors or DMSO control. At 9 hpi, the infectious virus particle production was assessed using the plaque assay. Data are presented as mean±SEM from at least three independent experiments. One-way ANOVA was used to evaluate statistical differences from the non-treated control and was presented when p≤0.05 with *p ≤ 0.05, **p ≤ 0.01, ***p ≤ 0.001, ****p ≤ 0.001.

### Hsp70 inhibition does not affect the number of intracellular vRNA copies

We next sought to identify the specific step of CHIKV replication most impacted by Hsp70 inhibition. To increase the resolution of the data, we increased the multiplicity of infection (MOI) from 1 to 10, thereby ensuring high infection rates. Before proceeding, we confirmed that both JG98 and JG345 are still able to significantly reduce the infectious particle production at MOI 10 (**Fig. S1**). CHIKV relies on the translation of its non-structural proteins and the formation of complex replication structures for effective genome replication, processes that may be disrupted by Hsp70 inhibition. Consistent with our findings that Hsp70 likely does not act during the early steps in the CHIKV infectious cycle, production of total vRNA (E1; **Fig. 5A**) and genomic RNA (Nsp1; **Fig. 5B**), as well as subgenomic RNA levels (**Fig. 5C**), were unaffected by JG98, at concentrations that reduced infectious particle production by 2-45-fold (**Fig. S1)**.

**Figure 5:**
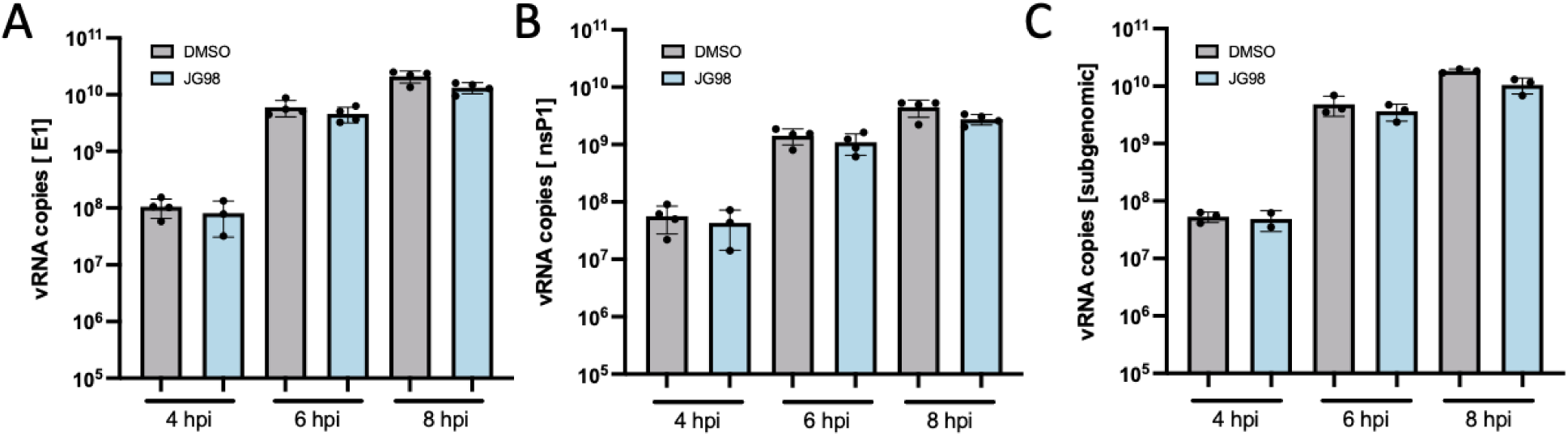
CHIKV genome replication and subgenomic vRNA formation are unaffected by Hsp70 inhibition. **(A)** The total intracellular vRNA and **(B)** genomic vRNA copy number were assessed at 4, 6, and 8 hpi in U2OS cells infected with CHIKV at MOI 10 and treated with vehicle control DMSO or 2μM JG98. Intracellular vRNA copies were quantified by RT-qPCR using specific primers against **(A)** E1 and **(B)** nsP1. **(C)** The number of subgenomic RNA copies was determined by subtracting the genomic vRNA copies from the total vRNA copies. Data is presented as mean ± SEM from at least three independent experiments. Student T-test was used to evaluate statistical differences from the DMSO control per timepoint and were presented when p≤0.05 with *p ≤ 0.05, **p ≤ 0.01, ***p ≤ 0.001, ****p ≤ 0.001.

### Expression of CHIKV viral proteins is reduced by Hsp70 inhibitors

We next assessed the effect of Hsp70 inhibition on the later steps after entry, i.e., viral protein expression using flow cytometry (see **Fig. S2** for gating strategy). In non-treated samples, approximately 50% of the cells were positive for CHIKV capsid and E2, and the solvent DMSO did not influence the percentage of cells expressing these viral proteins (**Fig. 6A and C**). However, treatment with both JG98 and JG345 significantly reduced the number of E2- and capsid-positive cells (**Fig. 6A and C**), a reduction further confirmed by a significant decrease in the mean fluorescence intensity (MFI) of E2 (**Fig. 6B**), though the MFI of capsid remained unchanged (**Fig. 6D**). This decrease in viral protein expression was validated using western blot analysis that showed a significant reduction in the levels of E2 expression with both JG98 and JG345 (**Fig. 6E and G**). However, the expression of capsid was only marginally reduced (**Fig. 6F and H**). Overall, these results suggest that the proper functioning of the Hsp70 machinery is crucial for the efficient production, correct folding, and/or stability of CHIKV proteins. However, since the reduction in viral protein expression does not directly reflect the decrease in infectious particle production, we hypothesize that Hsp70 inhibition may have additional effects during the late stages of replication beyond merely affecting protein expression.

**Figure 6:**
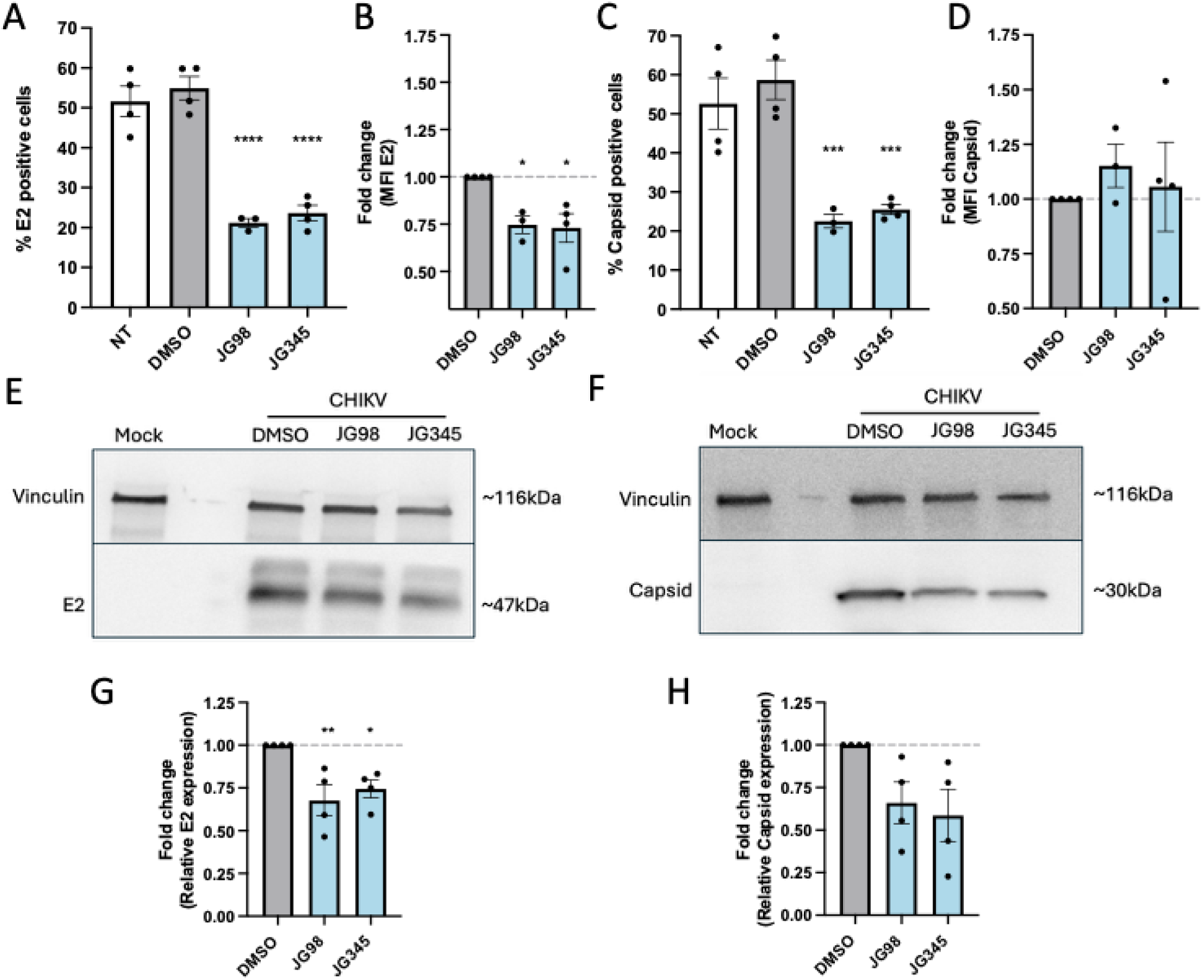
Viral protein expression is decreased by Hsp70 inhibitors. **(A-D)** U2OS cells were infected with CHIKV at MOI 10 and treated with JG98 (2μM), JG345 (1μM), or the DMSO control. At 9hpi, E2 and capsid expression were assessed using flow cytometry to determine the **(A and C)** percentage of positive cells and **(B and D)** mean fluorescent intensity (MFI; geometric mean). **(E and F)** Representative western blot of capsid, E2, or vinculin expression from protein lysates of U2OS cells infected with CHIKV at MOI 10 and treated for 9 h with Hsp70 inhibitor JG-98 (2μM), JG-345 (1μM), or the DMSO control. **(G and H)** Quantification of Western blots from three independent experiments. Protein levels are normalized to vinculin and are expressed as relative protein level to vehicle control DMSO. Data are presented as mean±SEM from at least three independent experiments. One-way ANOVA was used to evaluate statistical differences from the DMSO control and was presented when p≤0.05 with *p ≤ 0.05, **p ≤ 0.01, ***p ≤ 0.001, ****p ≤ 0.001.

### Hsp70 inhibition impairs the production of infectious CHIKV progeny

To test whether the Hsp70 inhibitors also affect later steps of the CHIKV replication cycle, including processes such as assembly and/or budding, we assessed the specific infectivity of particles produced with or without Hsp70 inhibitors present during infection with the CHIKV-LR strain. As demonstrated in **Fig. 1** and **Fig. 2**, Hsp70 inhibitors significantly reduced the number of infectious CHIKV particles produced (**Fig. 7A**). Indeed, the number of secreted genome equivalent copies (GECs) in the supernatant was significantly reduced by both JG98 and JG345 (**Fig. 7B**). To determine whether the compounds also may have affected the infectious potential of these particles, we calculated the PFU-to-GEC ratio, representing the amount of produced secreted genomes that are also infectious virions. The specific infectivity of produced CHIKV particles was reduced by at least 10-fold upon treatment with JG98 and JG345 (**Fig. 7C)**. The reduced PFU-to-GEC ratio reflects that Hsp70 inhibition also strongly affects the infectious potential of the virion rather than only the overall particle production. These data were confirmed using the CHIKV-S27 strain (**Fig. 7D-F**). Moreover, to determine whether the observed effects on progeny infectivity stem from disruption of Hsp70–co–chaperone interactions, we also assessed the impact of VER-155008 on CHIKV progeny production. Unlike the JG-compounds, VER-155008 inhibits Hsp70 by competing with ATP binding[49], thereby targeting this molecular chaperone machinery through a distinct mechanism. In comparison to the previous experiments (**Fig. 7A-C**), both the infectious (**Fig. 7G**) and total progeny production (**Fig. 7H**) were higher in both the NT and DMSO controls. Importantly, treatment with VER-155008 significantly reduced both the infectious (**Fig.7G**) and total progeny production (**Fig. 7H**) at concentrations that did not affect metabolic activity of the cells (**Fig. S3**). Notably, unlike the JG compounds, VER-155008 did not significantly impact the specific infectivity of CHIKV particles (**Fig. 7I**), suggesting that this effect is specific to the JG class of Hsp70 inhibitors.

**Figure 7:**
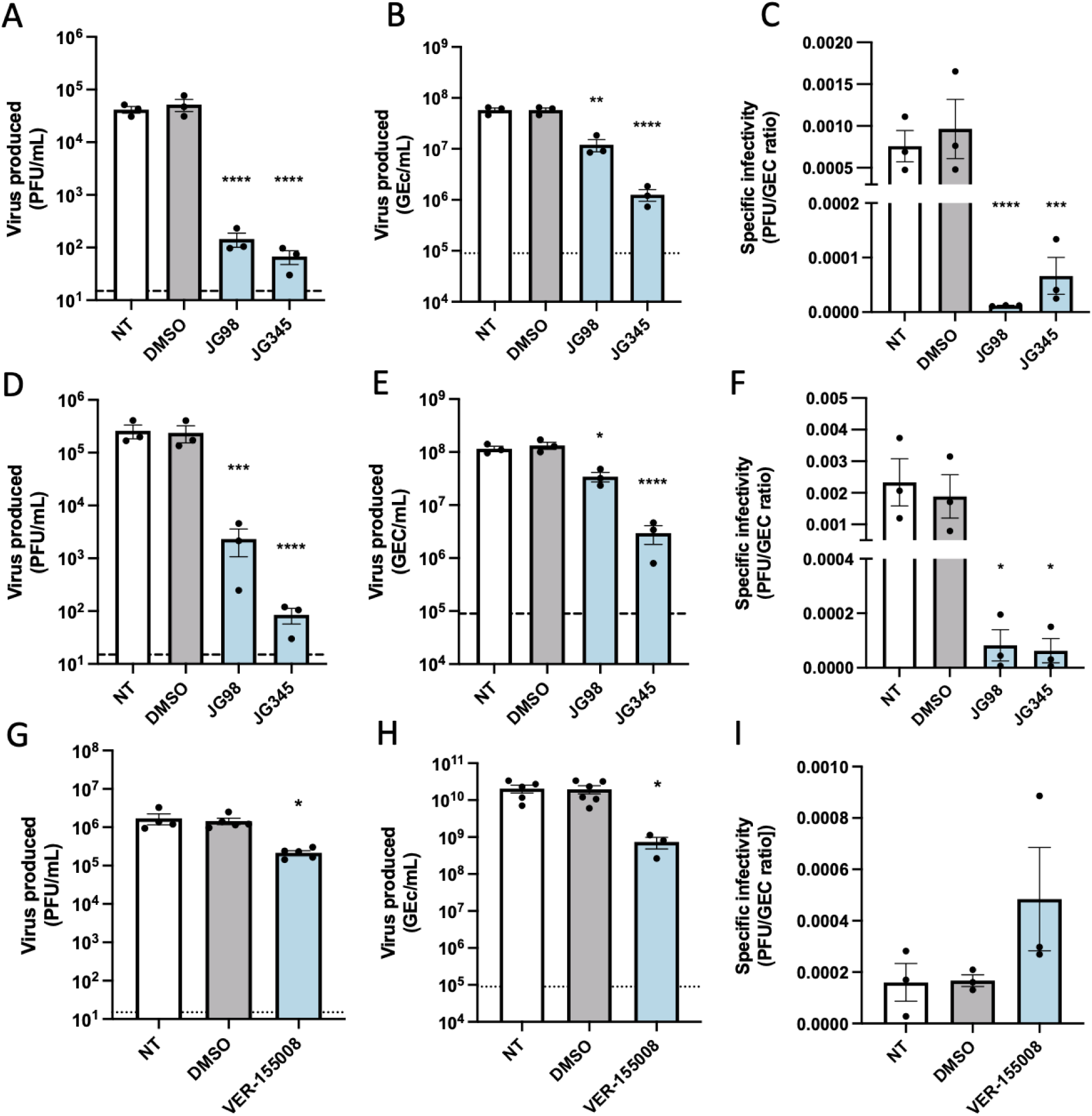
Hsp70 inhibition reduces the specific infectivity of produced CHIKV progeny. U2OS cells were infected with the **(A-C, G-I)** CHIKV-LR or the **(D-F)** CHIKV S27 strain for 9 h in the presence of **(A-F)** 2 µM JG98, 1 µM JG345, **(G-I)** 20 µM VER-155008, or the DMSO control. Supernatants were collected and the number of **(A, D, and G)** infectious particles and **(B, E, and H)** secreted genome equivalent copies (GECs) were measured using plaque assay and RT-qPCR, respectively. **(C, F, and I)** Specific infectivity is depicted as the ratio between produced infectious particles and GECs. Data are presented as mean± SEM from three independent experiments. One-way ANOVA was used to evaluate statistical differences from the DMSO control and was given when p≤0.05 with *p ≤ 0.05, **p ≤ 0.01, ***p ≤ 0.001, ****p ≤ 0.001.

### Inhibition of Hsp70 reduces ex vivo infection

To explore the antiviral potential of Hsp70 inhibitors in a more physiologically relevant setting, we next examined their effect in mouse skin explants, the initial site of CHIKV infection following a mosquito bite. We chose to use JG231 for this study, as studies showed that this compound has favorable pharmacokinetic properties over JG98 and JG345 in mice[31,33,34], and would therefore be most suitable for future *in vivo* studies. Before testing the compound in the skin explants, we first validated the antiviral activity in our model of U2OS cells and HFF-1 cells. To this end, cells were infected with CHIKV-LR in the presence of 2 µM JG231 or DMSO as a control. At this concentration, JG231 exhibited no cytotoxicity in either cell line (**Fig. S3**). Importantly, JG231 treatment significantly reduced the number of infectious viral particles in the supernatant of both U2OS and HFF-1 cells (**Fig. 8A and B**). Next, we assessed the *ex vivo* antiviral effect of JG231 in mouse skin explants infected with the CHIKV-899 strain, a clinical isolate (Mauritius, 2006) from the same lineage (ECSA) as CHIKV-LR, as previously described[50], during treatment with 2 µM JG231 or DMSO. In presence of JG231, the infectious titer (TCID₅₀ per mg tissue) was close to the limit of detection and 2.5-log lower than the DMSO control yet given the large variation in the DMSO control no statistical significance was reached (**Fig. 8C**). More constant results were obtained for the number of produced viral genome equivalent copies (GECs) per mg tissue and here a statistically significant reduction in virus particle production was detected compared to DMSO control (**Fig. 8D**). Together, these results demonstrated that Hsp70 inhibition in the skin using JG231 reduced the number of secreted CHIKV genome equivalent particles in the supernatant, thereby further substantiating the antiviral properties of JG-compounds.

**Figure 8:**
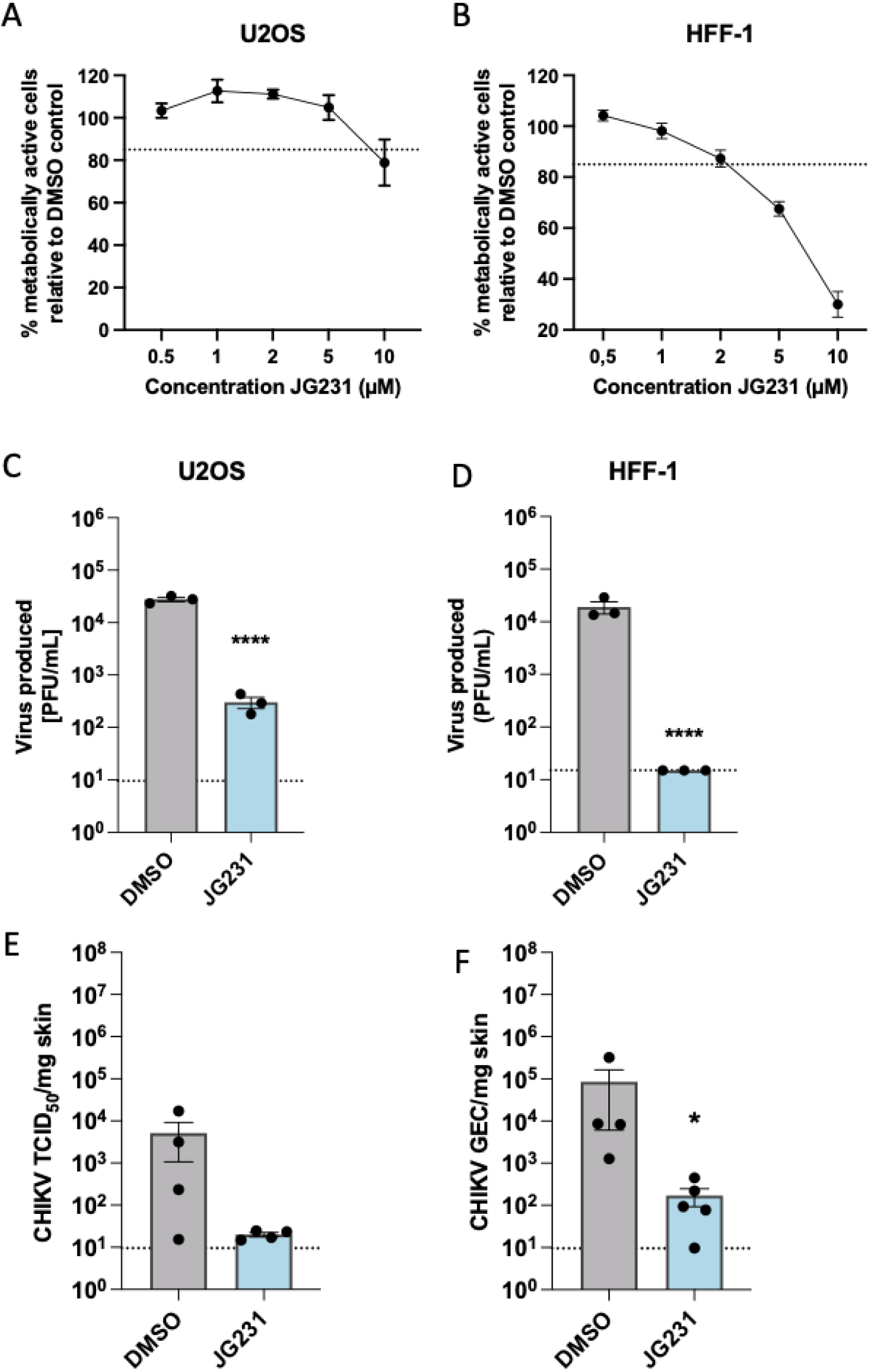
Inhibition of Hsp70 reduces CHIKV infection in *ex vivo* murine skin tissue. Metabolic activity of **(A)** U2OS and **(B)** HFF-1 cells was assessed after 16 h of treatment with increasing concentrations of JG231 or the equivalent volume of the DMSO control. The percentage of metabolically active cells is presented relative to the DMSO control. A decrease of 15% in metabolically active cells was considered non-toxic (dotted line represents 85%). **(C)** U2OS cells and **(D)** HFF-1 cells were infected with CHIKV-LR for 9 h in the presence of 2 µM JG231 or the DMSO control. Supernatants were collected, and the number of infectious particles in the supernatant was determined using the plaque assay. **(E and F)** Mouse skin explants were infected with 10^6^ PFU/mL of the CHIKV 899 strain during treatment with 2 µM JG231 or an equal volume of DMSO. **(E)** Infectious virus production (tissue culture infectious dose 50% (TCID_50_)) and **(F)** total virus production (GEC) at 2 dpi is displayed per mg tissue. The dotted line indicates the limit of quantification (LOQ). To evaluate statistical differences from the DMSO control, we used a Student T-test, and differences are presented when p≤0.05 with *p ≤ 0.05, **p ≤ 0.01, ***p ≤ 0.001, ****p ≤ 0.001.

## Discussion

The significant burden of the re-emerging arbovirus CHIKV underscores the urgent need for effective antiviral therapies, which are currently lacking. In this study, we investigated the role of the Hsp70 molecular chaperone in CHIKV replication and explored its potential as a target for antiviral strategies against CHIKV. Our findings reveal that allosteric inhibitors of the Hsp70-co-chaperone interaction (JG-compounds) effectively impaired the replication of multiple CHIKV strains across various cell types and in skin explants, with negligible cytotoxicity. While Hsp70 inhibition did not affect viral RNA replication, it significantly impacted the late stages of CHIKV replication, particularly protein expression. Furthermore, the decreased infectivity of CHIKV progeny produced under Hsp70 inhibition suggested that Hsp70 is crucial for additional late-stage replication processes, which might include processes like particle assembly and budding.

The involvement of the Hsp70 network in genome replication has been reported for several viruses, including DENV[36], ZIKV[37], Hepatitis C virus (HCV)[51], and Japanese Encephalitis virus (JEV)[52], typically through interactions with components of the viral replication complex (RC). Interestingly, we found that CHIKV genome replication and subgenomic RNA formation remained stable during treatment with Hsp70 inhibitors, suggesting that CHIKV RC formation is not dependent on Hsp70. One possible explanation for this Hsp70 independence is that CHIKV genome replication occurs within spherules at the plasma membrane[18,19], while flaviviruses like DENV, ZIKV, HCV, and JEV replicate their RNA in RCs located near the ER[53,54]. Due to differences in viral protein structure and localization, CHIKV may rely on different molecular chaperones than flaviviruses for RC formation. For instance, Heat shock 90kDa proteins (Hsp90s) have been proposed to aid in CHIKV RNA replication via interactions with components of the viral RC[55–57]. However, further research is needed to fully elucidate the distinct dependencies of RC formation on host molecular chaperones.

Although Hsp70 inhibitors (JG-compounds) did not affect CHIKV genome replication, they significantly reduced CHIKV capsid protein expression and the infectivity of progeny virions *in vitro*. Notably, however, no reduction in specific infectivity was observed in our *ex vivo* model with JG compounds, nor *in vitro* with VER-155008, an Hsp70 inhibitor that acts by competing with ATP binding. Importantly, in all conditions, there was a marked decrease in overall CHIKV particle production. Further research is needed to identify the exact mode of action of the compounds, yet it is tempting to speculate that Hsp70 may play a key role in the production and assembly of functional CHIKV capsids. Hsp70 is known to be involved in virion assembly for other viruses, such as HCV[58] and SV40[59]. While HCV capsids assemble in the ER[60] and SV40 in the nucleus[61], CHIKV assembles in the cytoplasm, likely relying on different Hsp70 homologs (and/or co-chaperones) for its assembly process. Late in infection, CHIKV also forms cytopathic vacuoles type-II (CPV-II), along with host vesicles, viral capsids, and glycoproteins[1], structures thought to contribute to capsid assembly[62]. Hsp70 may aid in these processes, and its inhibition could result in misassembled nucleocapsids due to defects in capsid-capsid or capsid-vRNA interactions. These defects can potentially compromise the stability or function of incoming vRNA during entry, thereby reducing the infectivity of virus progeny[63,64]. However, to pinpoint the exact role of Hsp70 in capsid assembly and/or CPV-II formation, more detailed studies on these processes during Hsp70 inhibition are needed.

Furthermore, JG98 and JG345 exhibited stronger inhibitory effects on CHIKV than JG18 and JG40. This finding was specific to CHIKV inhibition, as both JG18 and JG40 strongly reduced DENV replication. JG98, JG345, and JG231 belong to the type-2 JG-compounds, which are later-stage MKT-077 analogs with improved safety and potency compared to the type-1 compounds (e.g., JG18 and JG40)[34]. JG231 has even better pharmacokinetic properties than JG98 and JG345 and showed strong *in vitro* and *ex vivo* antiviral potential against CHIKV. Type-2 inhibitors have been shown to localize in the ER and vesicle-like structures[34], suggesting they might more strongly inhibit the ER-resident Hsp70 machinery, including HspA5 (BiP) and its specific co-chaperones (JDPs and NEFs)[65]. In this study, we found that Hsp70 inhibition significantly reduced the expression of the viral glycoprotein E2, which is processed in the ER, making it tempting to speculate that the ER-resident Hsp70 machinery indeed aids in CHIKV infection. The involvement of the ER-resident Hsp70 network has been well described in other viral infections, such as Simian Virus 40 (SV40)[35], where ER-resident JDPs (DNAJB11 and DNAJB12) and the ER-NEF (Glucose-regulated protein (GRP-)170) regulate viral exit from the ER into the cytoplasm[66]. Therefore, the role of JG-compound partitioning into cellular organelles during CHIKV infection should be further explored, as this may illuminate the importance of specific Hsp70 (co-)chaperones in CHIKV replication. Moreover, identifying CHIKV’s dependence on specific components of the molecular chaperone network could help refine targeting strategies and improve their tolerability[34].

This study highlights the critical role of Hsp70 in CHIKV replication and reinforces the existing evidence for the potential of targeting this chaperone machinery as an antiviral strategy against various arboviruses. However, given the varying antiviral efficacy of JG-compounds across different viruses, it is essential to carefully select the appropriate compound for each specific virus or identify a compound that has consistently strong antiviral effects towards distinct viruses. We show here that *ex vivo* inhibition of Hsp70 can reduce CHIKV replication in the skin, offering potential opportunities for localized treatment. However, before these compounds can be considered for clinical use, the *in vivo* antiviral potency against CHIKV must be thoroughly evaluated, and their pharmacokinetic properties further optimized[31,33]. Continued research to deepen our understanding of Hsp70’s role in CHIKV infection and to enhance the targeting of this host molecular chaperone is crucial for ultimately reducing the burden of CHIKV disease.

## Acknowledgments

We thank Dr. Izabela Rodenhuis-Zybert for her valuable input during work discussions.

## Supporting information

**Figure S1:**
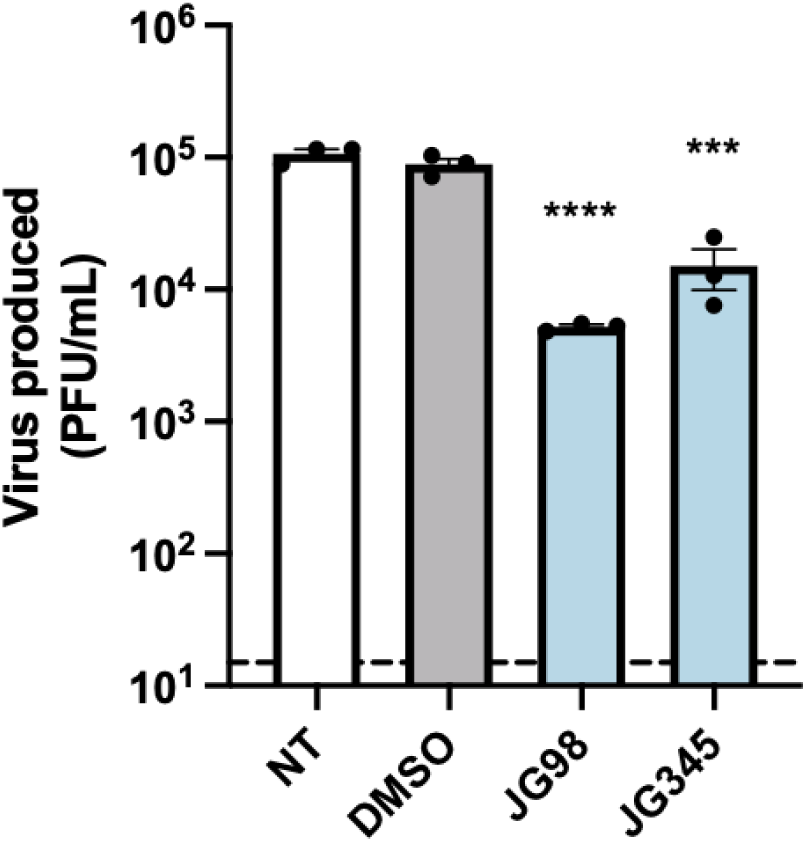
Progeny production of CHIKV infection at MOI 10 is still reduced by Hsp70 inhibitors. U2OS cells were treated with 2 µM JG98, 1 µM JG345, or the DMSO control during CHIKV infection at MOI 10. Virus production was measured at 9 hpi using a plaque assay. Data are presented as mean±SEM from at least three independent experiments. One-way ANOVA was used to evaluate statistical differences from the DMSO control and was presented when p≤0.05 with *p ≤ 0.05, **p ≤ 0.01, ***p ≤ 0.001, ****p ≤ 0.001.

**Figure S2:**
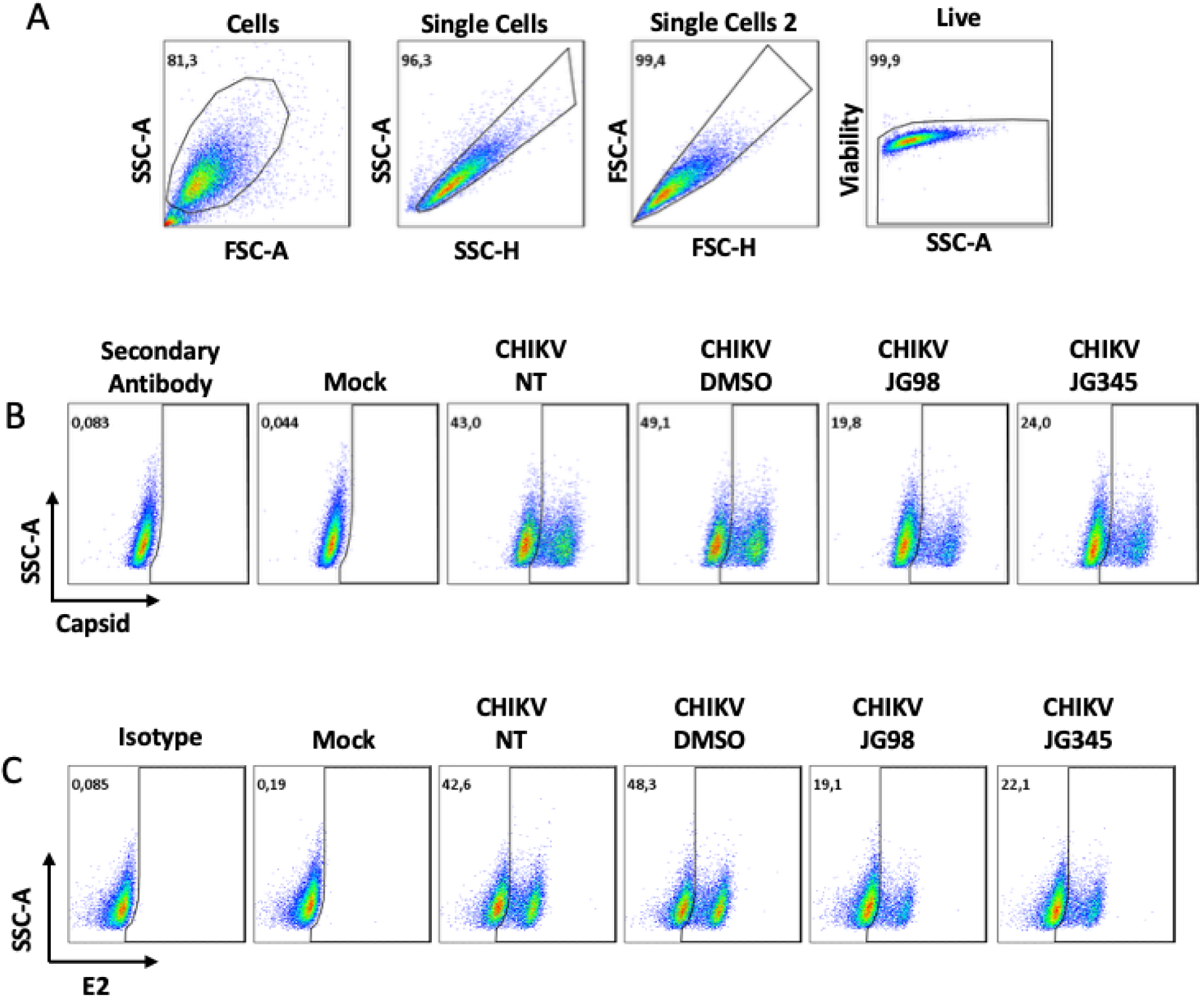
Gating strategy for flow cytometric analysis of CHIKV protein expression. Gating strategy to determine the percentage of cells positive for CHIKV E2 and capsid expression in U2OS cells treated with JG98, JG345, or the DMSO control. **(A)** Gating for cells and exclusion of doublets. **(B and C)** Gating to determine the percentage of cells positive for **(B)** capsid and **(C)** E2 based on mock-infected cells.

**Figure S3:**
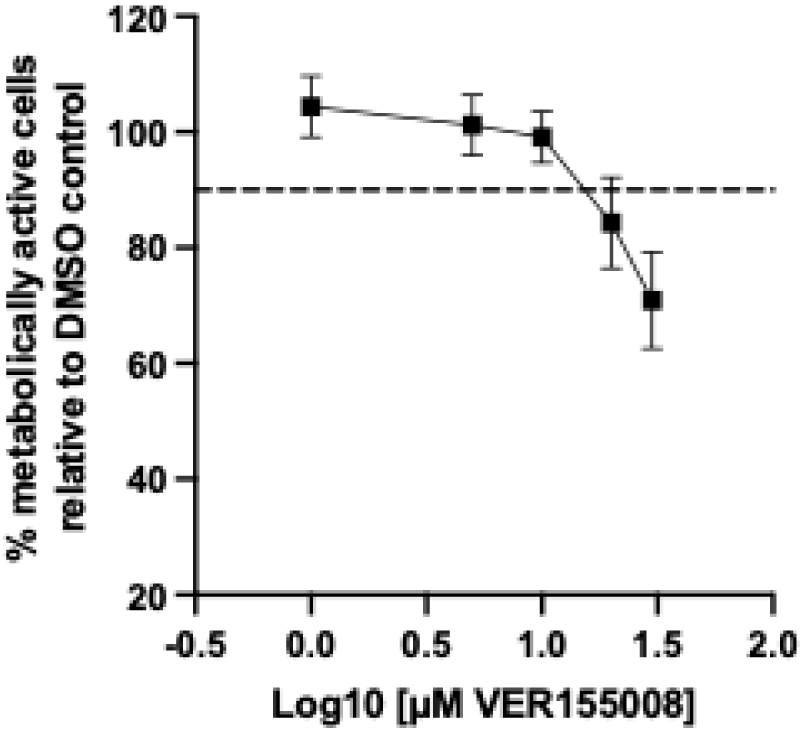
Dose-dependent effect of VER15008 on metabolic activity of U2OS cells. Metabolic activity of the U2OS cells was assessed after 16 hours of treatment with increasing concentrations of the VER155008 or the equivalent volume of the DMSO control. The percentage of metabolically active cells relative to the DMSO control. A decrease of 15% in metabolically active cells was considered non-toxic (dotted line represents 85%). Data are presented as mean±SEM from three independent experiments.

